# Antiviral action of aqueous extracts of propolis from *Scaptotrigona aff. postica* (Hymenoptera; Apidae) against Zica, Chikungunya, and Mayaro virus

**DOI:** 10.1101/2022.11.03.515030

**Authors:** RZ Mendonça, RM Nascimento, ACO Fernandes, PI Silva Junior

**Affiliations:** Laboratory of Parasitology, Butantan Institute; Laboratory for Applied Toxinology (LETA), Center of Toxins, Immune-response and cell signaling (CeTICS/CEPID), Butantan Institute

**Keywords:** *Scaptotrigona* aff *postica*, propolis, antiviral activity, Zicavirus, Chikungunya, Mayaro

## Abstract

The limited availability of antivirals for new highly pathogenic strains of virus has become a serious public health problem that kills thousands of people annually. For this reason, the search for new products against these agents has become an urgent necessity. Many studies have been carried out with this aim. Among the multiple sources of research for new antibiotics and antivirals, bioprospecting from insect exudates or their products has become an increasingly frequent option. Insects appeared on the planet about 350 million years ago and have been one of the beings with greater adaptability and resistance to the most varied biomes. Insects have been found in all known ecosystems. Their survival for so long, in such different environments, is an indication that they have a very efficient protection system against environmental infections, despite not having a developed immune system like mammals. Historically, since the ancient civilizations of Egypt and Rome, the products obtained from the bee, such as honey and propolis, have been of great pharmacological importance, being used as antimicrobial, anti-inflammatory, antitumor, healing several other functions. Investigations on the chemical composition and biological activity of propolis have been carried out, mainly in the species Apis mellifera, and this product has shown activity against some important viruses, such as poliovirus, influenza, HIV, hepatitis, and others. However, for the Meliponini species, known as stingless bees, there are few studies, either on their chemical composition or on their biological activities. The importance of studying these bees is because they come from regions with native forests, and therefore with many species of plants not yet studied, in addition to which they are regions still free of pesticides, which guarantees a greater fidelity of the obtained data. Previous studies by our group with crude hydroalcoholic extract of propolis demonstrated an intense antiviral activity against Herpes, influenza, and rubella viruses. All studies carried out with propolis are done with alcoholic extracts. In this work, we chose to use aqueous extracts, which eliminates the presence of other compounds besides those originally present in propolis, in addition to extracting substances different from those obtained in alcoholic extracts, which makes this work unprecedented. Therefore, this study aimed to identify, isolate and characterize compounds with antiviral effects from aqueous propolis extracts from *Scaptotrigona* aff *postica*, in emerging viruses such as zicavirus, chikungunya, and Mayaro. The evaluation of the antiviral activity of the crude and purified material was performed by reducing infectious foci in cultures of infected cells treated with propolis extracts in infected cultures and treated or not treated with propolis. The chemical characterization of the elements present in the extracts was performed by high pressure liquid chromatography. The results obtained indicate a high reduction of Zicavirus (64x) and Mayaro (256x) when was used 10% v/v of propolis and 256 x for chikungunya virus when was used 5% v/v of propolis. When compared to infected-only cultures. Even when was used 2% v/v of propolis, was observed a reduction of 128 fold in Mayaro virus replication. When purified fraction was used, the reduction observed was of 16 fold for Zicavirus, 32 fold for Mayaro virus and 125 fold for chikungunya virus. Likewise, it was observed that the antiviral response was dose-dependent, being more intense when propolis was added 2 hours after the viral infection. At the moment we are carrying out the chemical characterization of the purified compounds that showed antiviral action.

## 1. INTRODUCTION

Arboviruses are viruses transmitted by the bite of blood-sucking arthropods, such as Aedes aegypti. Aedes aegypti is a vector of great importance for public health in the tropics and subtropics and practically the entire American continent, as well as in Southeast Asia and throughout India. More than 200 arbovirus species of arbovirus have been isolated in Brazil, being around them related to diseases in humans (1). Arboviruses are emergent viruses by nature. None of them are originally humans disease. They are originally zoonoses, diseases of animals that only occasionally affect humans, and only become important when some significant ecological modification alters their natural habitat and leads to changes in reservoirs, vectors, and even virulence (2). The process of economic development, population growth and urbanization verified in Brazil from the second half of the 20th century placed arboviruses among the viruses of great impact, real or potential, on public health. In some situations, the adaptation to humans, of the virus or the arthropod vector, is sufficient to make the animal reservoir unnecessary for the maintenance of the virus cycle in nature. The main arthropod vectors are mosquitoes, flies, and ticks. The most common vertebrate hosts are rodents, but it also occurs in small mammals, primates, birds, and ungulates (3). In Brazil, the best-known and most widely circulated arboviruses with epidemiological importance today are Dengue, Zika, and Chikungunya.

Among the known arboviruses, Zika has left the world on alert since the recent outbreaks associated with it, especially in Brazil between 2013 and 2014, mainly in the Northeast region, with a high number of cases of microcephaly in newborns and other complications, as is the case of Guillain-Barré syndrome, which had not been reported in the same epidemiological proportion before. Although the Zika virus has been identified in humans in Africa in 1952 (4), the first major outbreak occurred only in 2007, in Micronesia (5). 2013 and 2014, outbreaks occurred in French Polynesia in 2013 (6) and New Caledonia in 2014 (7). A few times after this, still in 2014, reported cases of an illness characterized by rash and fever in northeastern Brazil. In May 2015, local transmission of the Zika virus is confirmed. In 2016, the Brazilian government reported 214,193 probable cases and 128,266 confirmed cases of Zika virus (8) although many cases were probably not reported. In this time is estimated that there were between 500,000 and 1.5 million cases from 2015 to early 2016 (9). As of May 2017, 85 countries and territories have reported documented cases of Zika virus transmission.

Recently, the Mayaro virus has become a potential public health risk for which there is no specific treatment. Less common in humans than Zica and Chikungunya, the Mayaro virus also causes outbreaks of febrile illness in the North and Midwest regions of Brazil, as well as in other South American countries (Bolivia, Peru, and Venezuela) (10). It belongs to the genus Alphavirus, family Togaviridae, along with Venezuelan equine encephalitis, eastern equine encephalitis, and Chikungunya viruses. Illness caused by the Mayaro virus is usually benign but can be temporarily disabling, often causing rash and arthralgia or even arthritis, myalgia, and fever lasting three to five days. During convalescence, arthralgia often persists for a few weeks. The sylvatic cycle of the Mayaro virus is similar to that of sylvatic yellow fever, a disease of monkeys, transmitted by mosquitoes of the genus Haemagogus. In addition to monkeys, the Mayaro virus can also be found in birds. All of them have a short life in the body, causing acute infections. In 2016, there was an explosion of cases of Chikungunya (63.810 confirmed cases) (11). There is a potential risk of the Mayaro virus becoming a virus with a greater public health impact than the previous two. The co-circulation of arboviruses in Brazil has made clinical diagnosis difficult, increasing cases of complications such as neurological fever syndromes, hemorrhages, and autoimmune diseases. The association of public policies for the prevention and control of vectors with advances in the development of vaccines capable of preventing the population against these diseases, maybe the solution for the control and reduction of cases of arboviruses in Brazil and the world. Another important aspect is the development of drugs that can inhibit the replication of the virus in the body and thus reduce the severity of clinical conditions. However, the importance of arboviruses is not concentrated only in Brazil. Recently, (12), the World Health Organization announce the launch of the Global Arbovirus. Arthropod-Borne viruses (Arboviruses) affect about approximately 3.9 billion people live health threats in tropical and sub-tropical areas. The frequency and magnitude of outbreaks of these arboviruses, particularly those transmitted by *Aedes* mosquitoes. Only dengue affect 89 contries. The Global Arbovirus Initiative is an integrated strategic plan to tackle emerging and re-emerging arboviruses with epidemic and pandemic potential focusing on monitoring risk, pandemic prevention, preparedness, detection and response, and building a coalition of partners. The initiative is a collaborative effort between the World Health Emergencies Programme, the Department of Control of Neglected Tropical Diseases, and the Immunization, Vaccines and Biologicals Department. This integrated initiative will build a coalition of key partners to strengthen the coordination, communication, capacity-building, research, preparedness, and response necessary to mitigate the growing risk of epidemics due to these diseases (12).

Despite the great importance in public health, few drugs are available for the treatment of these viruses after they are installed in the organism. Most treatments are only to overcome the effects of the virus and not to fight it. Seeking to combat these infections, the population has used natural products obtained from plants with antiviral actions for millennia. One of the products frequently used is propolis, a bee product that has a very wide range of pharmacological actions. Among the best-known activities of propolis, its antiviral action is one of the most outstanding, as it demonstrates better efficacy against viruses than standard drugs available on the market (13). An example of this is the comparative study between an ointment prepared with Canadian propolis, rich in flavonoids, and acyclovir ointment (a drug indicated for the treatment of Herpes simplex, Varicella zoster, Epstein-Barr and Cytomegalovirus viruses). In this study, the propolis-based ointment was more effective than acyclovir in healing genital herpes lesions and in reducing local symptoms in the treatment of genital herpes simplex virus (14). An in vitro study with Canadian propolis proved to be efficient against Herpes simplex, types 1 and 2 during the viral adsorption phase or when incubated directly with the virus. These results indicate that propolis can directly interfere with the virus, preventing the penetration of the virus into cells (15). Propolis harvested in the city of Moravia, the Czech Republic, when tested against HVS-1, showed a protective effect when the virus was previously incubated with the extracts (16). The composition of the word propolis comes from the Greek, pro means “in defense”, and polis means “city”, that is, “in defense of the city”. Propolis is a resinous material that bees collect from plants, mix with wax, and use to build their nests. Bees apply propolis in thin layers to the inner walls to strengthen and block holes and crevices, as well as use it as an “embalming” substance on dead invaders or even dead bees, thus inhibiting the growth of fungi and bacteria inside the hive. (17, 18) The medicinal properties of propolis have been reported since ancient times. In Egypt it was used in the mummification process to prevent bodies from rotting; the Greeks and Romans used it as an antiseptic and healing agent, and the Incas as an antipyretic (18, 19) Due to its great pharmacological potential, propolis became the target of several researches, arousing the interest of the scientific society that has sought to investigate its compounds and biological activities. This interest has intensified in recent decades (17, 20). Countries such as Switzerland and Germany already legally recognize propolis as a medicine (21). In Brazil, the main substances described are those derived from prenylates of p-coumaric acid and acetophenone, such as diterpenes and lignans (21). Studies with propolis have already demonstrated biological activities, such as immunomodulation, antitumor, antioxidant, antibacterial, antiviral, and analgesic (22, 23, 24).

One of the components of hives is resins. Resins are pre-formed substances produced by secondary metabolites of plants, which have specific functions, such as pigmentation of flowers to attract pollinating insects, repellent action, and defense mechanisms against pathogenic microorganisms (25). In propolis, the resin is used in prophylactic protection and combat against microorganisms. Studies show that epidemics in nests decrease both productivity (26) and the number of bees (27). Therefore, the chemical compounds present in these products are essential for the protection and survival of bees, being considered an evolutionary mechanism for the benefit of the colony (28). The composition of propolis is extremely variable, both in terms of geographic location and season (29). In Europe, North America, and temperate zones, flavonoid compounds, phenolic acids, and esters predominate in propolis (30). In the Mediterranean terpenoids have been identified in high concentrations (31,32). In Africa, the main compounds are triterpenoids (33).

In Brazil, due to the great biodiversity, several types of propolis have been reported, such as red, green, and brown. The vast majority of studies are carried out with propolis from bees of commercial interest, mainly Apis mellifera. The propolis of the species Apis mellifera was classified into 12 distinct groups, based on the physicochemical characteristics and place of origin. Among the main compounds isolated are coumaric acid, ferulic acid, pinobanksin, kaempfero, apigenin, and caffeic acid (34). The components of Brazilian red and green propolis are commonly extracted from Baccharis spp, Dalbergia ecastaphyllum, Betula verrucosa, Betula pendula, and Betula pubescens species and are composed of phenylpropanoids, phenolic acids, p-coumaric acids, diterpenic acids (33). Studies on the glandular secretions of bees have demonstrated antibacterial activity, as is the case with the hypopharyngeal secretion of the species Apis mellifera (35). The same hypopharyngeal glands were found in the species *Scaptotrigona postica*, presuming they have the same functions (36). The vast majority of studies on the pharmacological effects of propolis are related to propolis obtained from Apis mellifera bees (19). The chemical composition of propolis is influenced not only by the vegetation of the environment but also by the seasons and the bees that produce it (17). These variations in chemical components affect their biological properties (37). For this reason, it is so important to study the propolis of different bee species and regions. Among the different species of bees, studies dealing with propolis from stingless bees are rare (38). In Brazil, one of the species of stingless bees belongs to the Apidae family, of the Meliponinae subfamily. These bees mix plant resin with wax and clay, thus forming a geopropolis. This denomination is precisely due to the use of land for the final composition of the material produced (38,39). Recently, compounds such as phenolic acids and water-soluble tannins were isolated from the geopropolis of Melipona fasciculata, (gallotannins and ellagitannins). These compounds, from the aqueous and alcoholic fractions of geopropolis from Melipona scutellaris (40,41), demonstrated antioxidant activity (42), anti-inflammatory and antinociceptive activities. Within the Meliponinae subfamily, there is the genus Scaptotrigna, which is distributed throughout the Neotropical region and includes species that build their nests in pre-existing cavities (43). In this genus, we have *Scaptotrigona* aff *postica*, popularly known in Maranhão, a region of northeastern Brazil, as the “tubi” bee. However, despite being in the family of geopropolis producers, they are an exception to the rule, since they do not use land in the composition of propolis (20).

There are few studies with propolis from *Scaptotrigona* aff *postica*, and there is no consensus in the literature on how to refer to it, some groups use the name geopropolis (46), while others use propolis (20,44, 45). For this work, we will use the term propolis, as we understand it to be the most appropriate. This propolis has been used by the population of the Maranhão region in the treatment of cancer and wound healing (20). Earlier, we have shown that propolis from *Scaptotrigona* aff *postica* presents a potent antiviral action against Herpes Simplex Virus (HSV-1) (46) as well against Rubella virus (47). Scientific studies have corroborated the possible medicinal effects of this propolis. Recently, we have shown (48) that propolis and its components can help reduce the physiopathological consequences of covid-19 infection, and that, flavonoids glycosides and pyrrolizidine alkaloids from propolis of *Scaptotrigona* aff. *postica* has a potent antimicrobial action (49). Martin et al. (50) treated rats with corneal injuries caused by burns with an emulsion of the crude extract and observed both healing and anti-inflammatory effects. Antitumor activity against Ehrlich tumor and reduction in pathology associated with asthma due to an inhibition of migration of inflammatory cells into the alveolar space are also described. In addition to those mentioned, there are no reports on other actions of this product. Therefore, due to the importance of these arboviruses in public health, the limited availability of effective drugs on the market, and the abundant information on the antiviral activity of propolis from several bee species, this work aims to study the antiviral activities of *Scaptotrigona* propolis. aff *postica* against arboviruses.

## 2. Material and methods

### 2.1 Propolis

The propolis used in this work was obtained from a colony of *Scaptotrigona* aff *postica*, located in the State of Maranhão, in the Barra do Corda region, Brazil from apiary Melinative. The city is located in the geographic center of Maranhão, at the confluence of the Rio da Corda and Rio Mearim.

### 2.2 Preparation of crude extract of propolis

In this work, the aqueous extraction of propolis was used as a source of substances for studies of antiviral action. Aqueous extraction of propolis was carried out as described below. The propolis, obtained by scraping the meliponiculture box, was gathered, forming a “paste”. This material was transported to the laboratory and frozen at −20°C for 24 hours to make the material rigid, forming a frozen propolis “stone” to facilitate the spraying process. This “stone” of propolis was then manually macerated until it became a very fine granular material. This crushed material was passed through stainless granulometric sieves, fine mesh of 0.15 mm. The sieved material was ground again by one degree until the material above became a very fine powder which was passed again through ABNT (Brazilian Association of Technical Standards) granulometric (75 to 100 mm Astn) sieves with a porosity of 1 to 5 microns. After this procedure, ultrapure water was added to the powder (100 ml of water for every 10 grams of propolis powders) and homogenized under rotation at 1,000 rpm for 24 hours at room temperature, protected from light. To separate the precipitate (wax) from the liquid medium (extract), the supernatant was centrifuged at 4°C for 30 minutes at 15,000 rpm (Eppendorf® centrifuge 5804R). The supernatant was then filtered through a 0.22µm millipore membrane, aliquoted, and stored in a sterile glass refrigerator until use. This material was considered to be 100%.

### 2.3. Cell line, culture medium, and cell culture

VERO cells (African green monkey kidney fibroblast cells - *Cercopithecus aethiops*) (Vero - CCL-81, obtained from the ATCC) were used in this study to determine the antiviral action of propolis. These cells were cultured at 37°C in T-flasks or 96-well microplates containing Leibovitz medium (L-15), supplemented with 10% fetal bovine serum. Cells were maintained at 37°C in a cell culture oven. Cell growth and morphology were monitored daily using an inverted optical microscope Olympus CK2 at 200x magnification.

### 2.4. Cytotoxicity assay of samples in cells

The cytotoxicity test of all samples and the purified fractions of propolis was performed in VERO cells to determine the maximum concentration of propolis or its fractions that is not toxic. For this, cells were seeded at a concentration of 10^4^ cells/well in 96-well plates and cultured at 37°C for 24 hours. After this period, the cells were treated with different concentrations of the samples to be tested. PBS was used as a negative control and DMSO 10% as a positive control. After 24, 48, and 72 hours of treatment, cell viability was determined by visualizing cell morphology and treating the cells with a vital dye, trypan blue. In each experiment, an aliquot of the samples, at the concentration to be used in the tests, was added to a plate excavation containing VERO cells, as a way of verifying the non-toxicity of the sample, serving as a basis for comparing the morphology of the cells in the test excavations.

### 2.5 Vírus and infection

#### 2.5.1 Virus

The antiviral action of *Scaptotrigona* aff *postica* propolis was tested in 3 different arbovirus of interest in public health; (Zicavirus, Chikungunya, and Mayaro). These virus are part of the collection of the Parasitology laboratory at Instituto Butantan

#### 2.5.2 Viral titulation

Initial viral stocks were titrated at the beginning of the work by the viral extinction technique as described by Reed and Muench. (51) and detailed in the following item. The viral titer was expressed as TCID50/mL (50% Tissue culture infectious dose), which is the highest dilution of the virus capable of infecting 50% of the cells. To determine the viral titer, aliquots of the virus samples under study were placed on a semi-confluency mat of VERO cells (2 x 10^5^ cells/mL), grown in 96 microplates. The viruses were added to the cell cultures in serial dilutions 2. The microplates were incubated at 37°C and the viral cytopathic effect was observed daily and the titer of infectious viral particles was calculated as described by Reed and Muench. For this study, an aliquot of the virus was thawed immediately before experimenting.

#### 2.5.3 Test of antiviral activity

##### 2.5.3.1. Test of antiviral activity in cell culture by the technique of determination of the minimum infective dose (TCID/50)

In the first phase of tests, VERO cells, an exponential growth phase, grown in a 96-well plate, were treated with crude samples of the test substances, 1 hour before infection. After this treatment period, the cells were infected with the different viral strains in serial dilutions ratio 2, with the first dilution varying from 100 to 500 TCI/50 of virus. The cultures were kept at 37°C and observed daily under an optical microscope, seeking to determine the inhibitory effect of the samples on virus replication by determining the cytopathic effect. After the appearance of the cytopathic effect, the culture supernatant was removed and the cells were stained with crystal violet to reveal the cytopathic effect. The antiviral effect was determined by the difference in the highest dilution of the virus capable of inducing a cytopathic effect observed in the infected-only culture and the infected but treated culture. Negative controls for infection and cytotoxicity of the samples were performed in all experiments.

##### 2.5.3.2 Dose/effect determination

To define the best amount of fraction to be added to infected cultures, fractions showing antiviral activity were added 1 hour before infection at concentrations of 2, 5, or 10% (v/v). The cultures were kept at 37°C and observed daily under an optical microscope, seeking to determine the inhibitory effect of the samples on virus replication through the determination of the cytopathic effect and the determination of the effect performed as described above.

##### 2.5.3.3 Determination of the optimal time of infection (TOI)

To define the best time of infection (TOI), the fractions that showed antiviral activity were tested for the best time to add these fractions to the infected cultures. For this, samples of the fractions, at the optimal concentration described above, were added 2 hours before infection, and 2 hours after infection, or the cultures were treated with a mixture of fraction + virus and kept in contact for 2 hours. before being added to crops. The cultures were kept at 37°C and observed daily under an optical microscope, seeking to determine the inhibitory effect of the samples on virus replication through the determination of the cytopathic effect and the determination of the effect performed as described above. Six days after the start of the experiment, the supernatant was removed, 0.25% Crystal Violet was added and washed with PBS

### 2.6. Fractionation of samples by Reverse-Phase HPLC and antiviral activity test of purified fractions

#### 2.6.1 Aqueous Propolis Extract HPLC Purification

*Scapotrigona aff, postica* propolis (1,2g) was suspended in 180 mL of ultrapure water and homogenized for 2 hours at 4°C. The supernatant (20 mL) was fractionated by a reverse-phase high-performance liquid chromatography (RP-HPLC) at room temperature on a Shimadzu LC-10 HPLC system using a semi-preparative C18 Jupiter column (30 mm; 300A; 10 LJ 250 mm) (Phenomenex International, Torrance, CA, USA) equilibrated at room temperature with 0.1% TFA in ultrapure water. The elution was done under a linear ACN gradient and the flow rate was 8 mL/min during 60 min (0-80% of B). Each fraction (1mL) was individually and manually collected, under Ultraviolet (74) absorbance monitored at 225 nm. Eluted fractions were immediately refrigerated, concentrated in a vacuum centrifuge SpeedVac Savant (Thermo Fisher Scientific, Waltham, MA, USA), and stored in the freezer at −20 °C until its usage. The concentrated fractions were reconstituted in 1 mL ultrapure water and used in antiviral activity assays. The purified fractions were tested for their antiviral capacity in the same way as performed with the crude propolis sample.

#### 2.6.2 Characterization of propolis fractions with activity by mass spectrometry

The substances with activity were resuspended in 20 μL in ACN/H2O 1:1 with 0.1% formic acid and analyzed by direct infusion in a Thermo Scientific LTQ-Orbitrap LC/MS (LTQ – Linear Trap Quadrupole), under 0.5 μL/min in a 50 μL syringe (Hamilton, Reno, NV, USA).

### 2.7 Protein dosage

The concentration of the fractions was determined by the Lambert-Beer law, using the molar extinction coefficient at 205 nm absorption (52). The active fractions concentration was by measuring A205 nm using a NanoDrop 2000c spectrophotometer (Thermo Fisher Scientific).

## 3 RESULTS

### 3.1. Determination of the cytotoxicity of the agents to be tested

At the beginning of the work, the propolis cytotoxicity of *Scaptotrigona* aff. *postica* was determined in VERO cells. The concentrations used were the same used in the tests. For this, VERO cells were grown in 96-well microplates. When the cell reached confluently, 1, 2, 5, or 10% v/v propolis or its active recombinant analog was added to the cultures in triplicate. The cultures were observed daily and for 72 hours to determine the morphological alteration in the cultures. None of the concentrations used showed cytotoxic effects in the cultures. As a negative control of the experiments, in all the tests performed, an aliquot of the agent to be tested was always added to the normal VERO cells, at the same concentration as the test and used for comparison with the results obtained. None of the concentrations tested had a deleterious effect on the growth of Vero cells (**Figure 01**). Cells were observed under an inverted microscope at 10x magnification.

**Figure 1:**
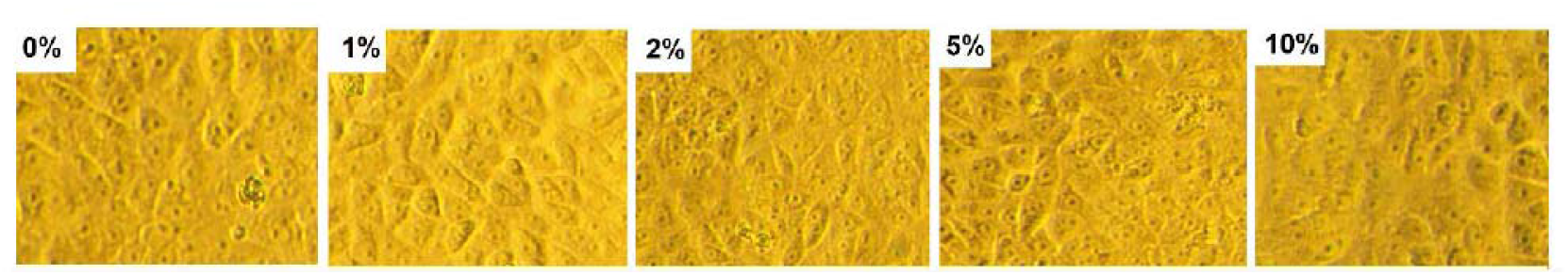
Cell viability of vero cells treated with different concentrations propolis. The culture was treated con 1, 2, 5 or 10% v/v of propolis. The culture was maintained at 37 °C for 72 hours and the culture was observed daily to determine the cytotoxic effect. The results above represents the morphology observed in cells in all experiments.

### 3.2 Chromatography of the propolis aqueous extract and identification of fractions with viral activity

To identify the molecules present in the sample with antiviral action, phytochemical analysis of the propolis extract was performed. The extract was fractionated by a reverse-phase high-performance liquid chromatography (RP-HPLC) at room temperature on a Shimadzu LC-10 HPLC system using a semi-preparative C18 Jupiter column (30 mm; 300A; 10 LJ 250 mm) (Phenomenex International, Torrance, CA, USA) equilibrated at room temperature with 0.1% TFA in ultrapure water. The elution was done under a linear ACN gradient and the flow rate was 8 mL/min during 60 min (0-80% of B) (**Figure 02A**). Each fraction (1mL) was individually and manually collected, under Ultraviolet (74) absorbance monitored at 225 nm. Eluted fractions were immediately refrigerated, concentrated in a vacuum centrifuge SpeedVac Savant (Thermo Fisher Scientific, Waltham, MA, USA), and stored in the freezer at −20 °C until its usage. The concentrated fractions were reconstituted in 1 mL ultrapure water and used in antiviral activity assays. The purified fractions were tested for their antiviral capacity in the same way as performed with the crude propolis sample. The fraction 1 (with antiviral activity) was subjected to mass spectrometry using a Thermo Scientific LTQ-Orbitrap LC/MS (LTQ – Linear Trap Quadrupole). As can be observed in **Figure 02B**, various iond were obtained. These ions were submitted to MagTran ® for mass analysis and identification (deconvolution), and revealed a molecule of mass of 1,331.96 Da and a concentration of 107µg/ml.

**Figure 2:**
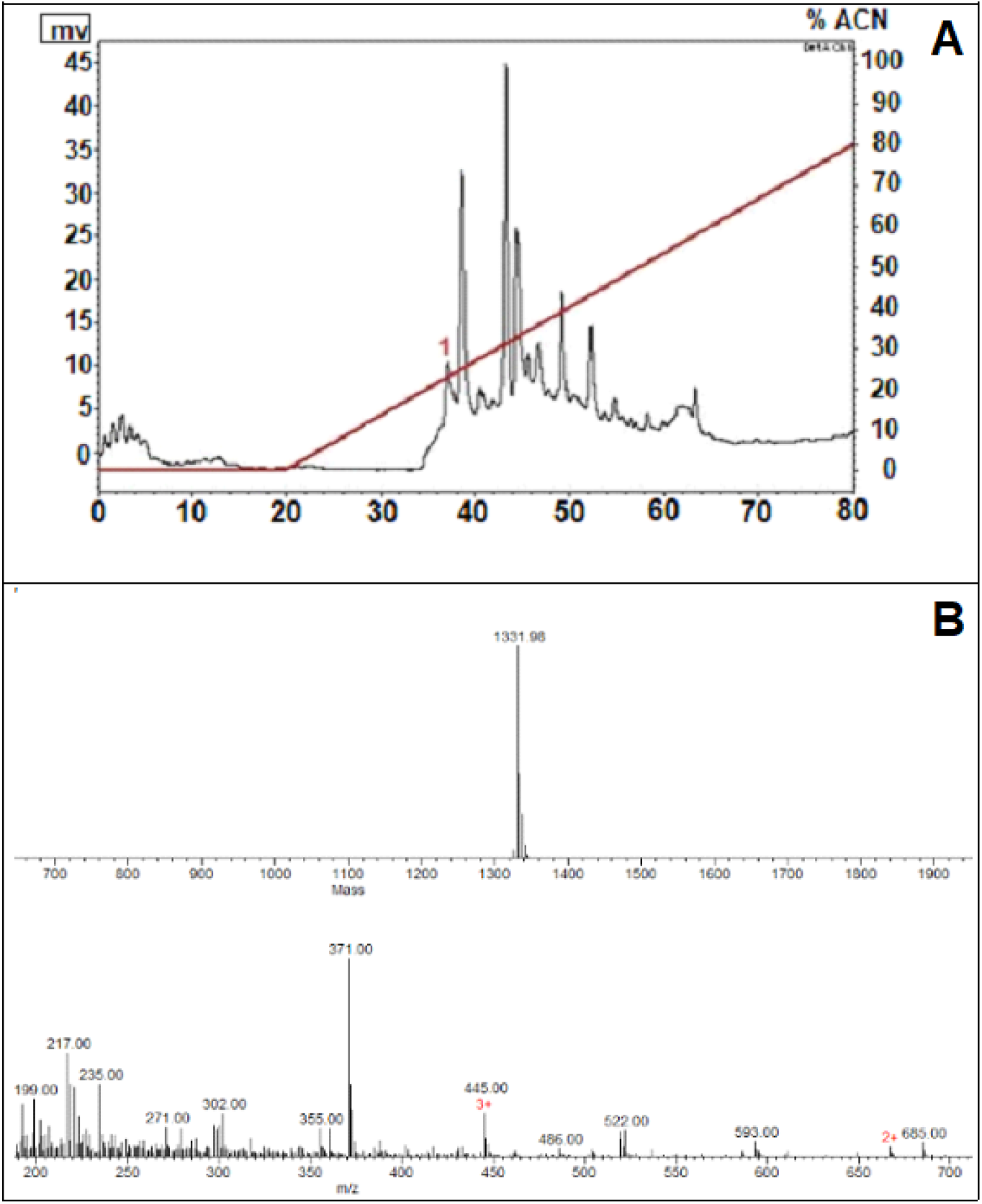
Phytochemical analysis of the propolis aqueous extract. **A**-Reverse-phase high-performance liquid chromatography (RP-HPLC): samples eluted with a linear gradient from 0% to 80% ACN in acidified water for 60 min (8 ml/min.). The fractions were collected manually and submitted to a test of the inhibition of microbial growth in a liquid medium. **B**- The fraction 1 (with antiviral activity) was resuspended in 20 μL in ACN/H2O 1:1 with 0.1% formic acid and analyzed by direct infusion in a Thermo Scientific LTQ-Orbitrap LC/MS (LTQ – Linear Trap Quadrupole), under 0.5 μL/min in a 50 μL syringe (Hamilton, Reno, NV, USA). Mass spectrometry analyses revealed some ions (m/z) the most abundant being an interferent (371 m/z). Ions were submitted to MagTran ® for mass analysis and identification (deconvolution), and revealed a molecule of mass of 1,331.96 Da.

### 3.3 Determination of antiviral action

#### 3.3.1. Determination of the antiviral action of propolis on different arboviruses

In this work, the antiviral action of propolis was tested against Zica virus, Chikungunya, and Mayaro virus. For this experiment, plates from 96 wells with VERO cells were treated with propolis being maintained in contact with the cells for 1 hour. After this period, serial dilutions (ratio 2) of the virus, with an initial titer of 100 to 500 TCID/50, were added to the treated cultures and observed for 72 hours to determine the cytopathic effect.

##### 3.3.1.1. Antiviral action of propolis and its purified fraction against Zica virus

The test was performed as described above. As can be seen in Figure 3, there is an intense reduction in the viral effect of 256 TCID/50 of Zica virus after the treatment of cells with 10% v/v propolis. Under these conditions, the viral reduction ranged from 256 to 4 TCID/50, or 64 fold (98,4%), with crude propolis. With the treatment of cells with purified fraction 1, the antiviral action was 16-fold (256 to 16 TCID/50).

**Figure 3:**
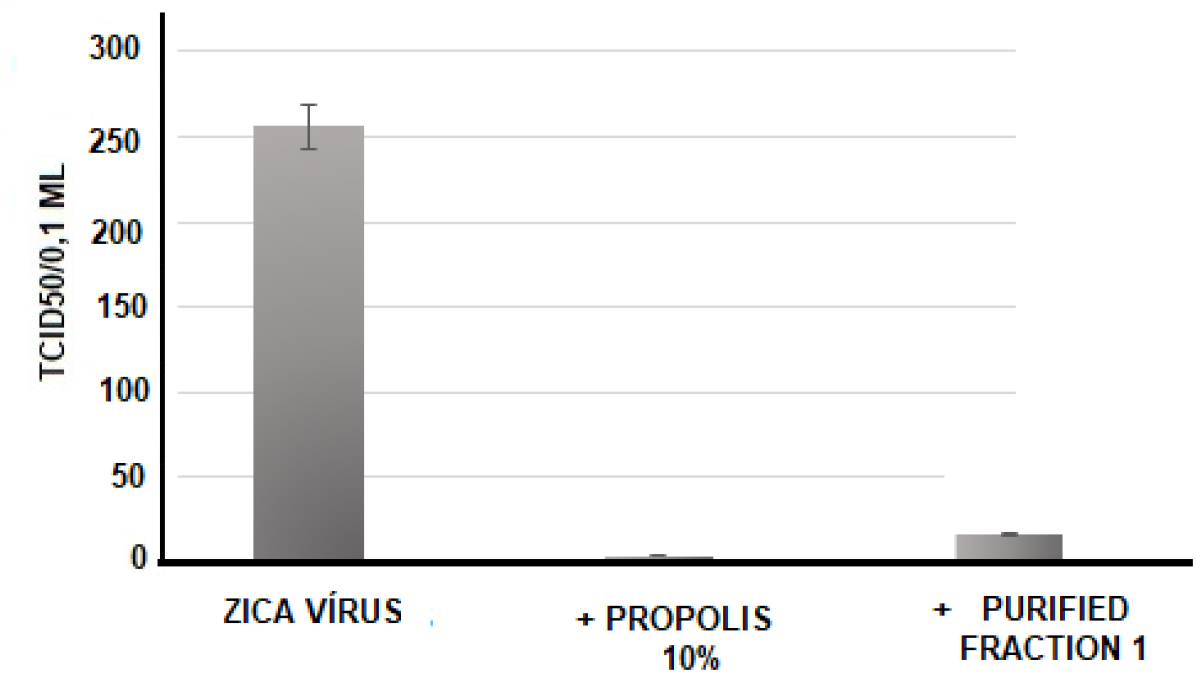
Antiviral action of propolis from *Scaptotrigona* aff. *postica* (10% v/v) against the Zica virus. The propolis extract or the purified fractions were added to the cultures of VERO cells and 1 hour later, serial dilutions (r:2) of Zica virus (initial titer of 256 TCID 50/0.1ml) were added to the culture. The culture was maintained at 37 °C for 72 hours and the culture was observed daily to determine the cytopathic effect. The final titer was determined by the highest dilution of the virus to cause a cytopathic effect in the culture. The result represents 4 individual experiments (performed in duplicate each).

In the photos in Figure 4, can be seen that there was a clear reduction in the cytopathic effect (B) caused by the Zicavirus 72 hours after infection, in cultures treated with propolis (10% v/v) (C), which presents a normal morphological aspect as control cells (A).

**Figure 04.**
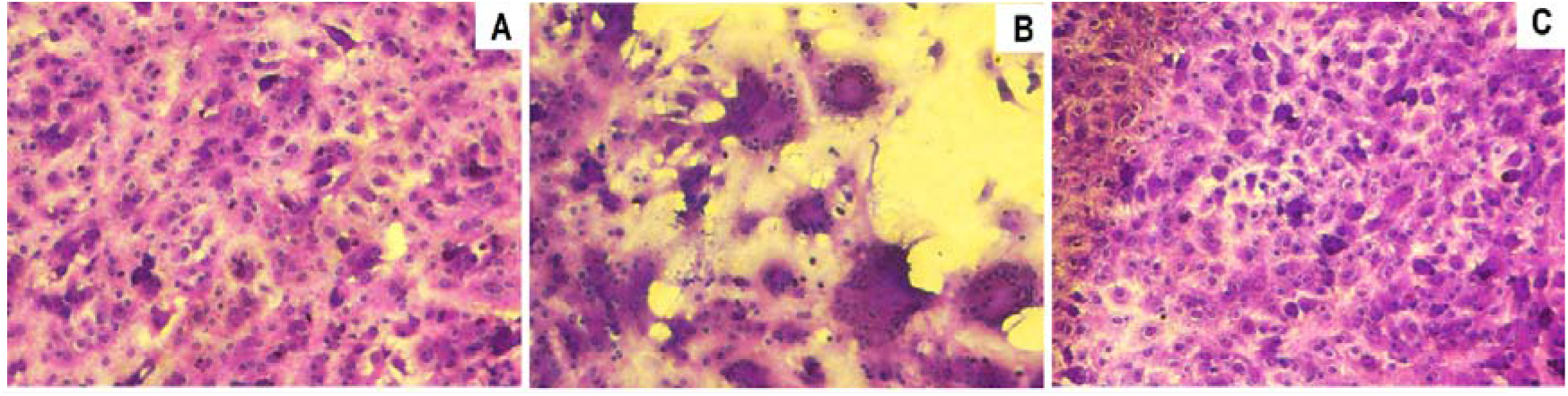
Photomicrography of normal VERO cell cultures (A), infected with Zica virus (B) and infected and treated with 10% v/v propolis from *Scaptotrigona* aff*. postica* (C). The cultures were kept for 3 days at 37°C. Cells were observed daily under an inverted microscope at 10x magnification. After this period, the culture supernatant was removed and the cells stained with crystal violet (0,25%).

##### 3.2.1.2 Antiviral action of propolis against Mayaro virus

The antiviral action of the aqueous extract of propolis was also tested against the Mayaro virus. The experimental procedures were the same as mentioned above but using 512 TCID/50 of virus and 10 % propolis. Under these conditions, the viral reduction was similar to that obtained in the tests with the Zica virus, using crude or purified propolis fraction. In this case, the reduction was on average 128-fold for crude propolis (512 to 4 TCID50) and 32-fold for the fraction (512 to 16 TCID/50) (Figure 5). The result represents 4 individual experiments (performed in duplicate each).

**Figure 5:**
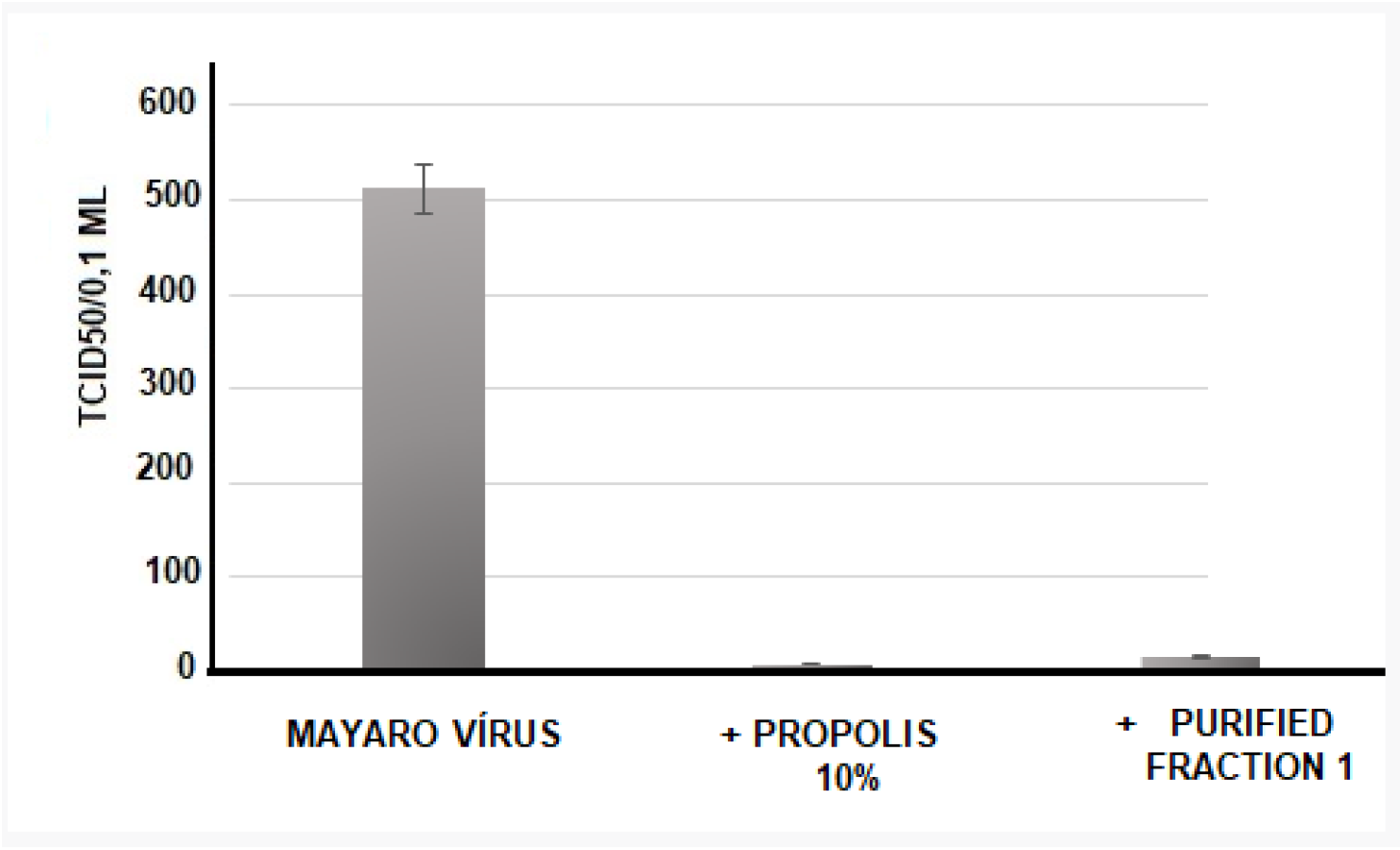
Antiviral action of propolis from *Scaptotrigona* aff. *postica* against the Mayaro virus. Aqueous propolis extract (10% v/v) was added to the VERO cell culture. One hour after this procedure, serial dilutions (r:2) of the Mayaro virus (512 TCID 50/0.1ml) were added to the culture. The culture was maintained at 37 °C for 72 hours and the culture was observed daily to determine the cytopathic effect. The final titer was determined by the highest dilution of the virus to cause a cytopathic effect in the culture.

In the photos in Figure 6, can be seen that there was a clear reduction in the cytopathic effect caused by the Mayaro virus 72 hours after infection, in cultures treated with propolis (2 % v/v).

**Figure 06.**
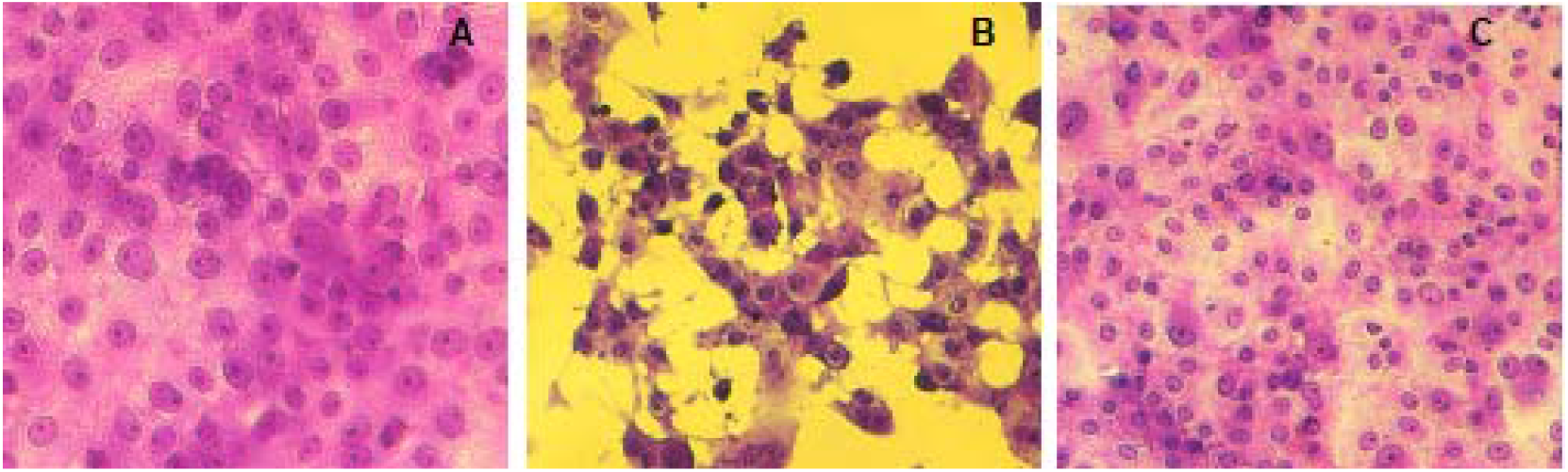
Photomicrography of normal VERO cell cultures (A), infected cells with Mayaro virus (B), and infected cells treated with 10 % v/v propolis from *Scaptotrigona* aff. *postica* (C). The cultures were kept for 3 days at 37°C. Cells were observed daily under an inverted microscope at 10x magnification. After this period, the culture supernatant was removed and the cells were stained with crystal violet 025%.

##### 3.2.1.3. Antiviral action of propolis on Chikungunya virus

The antiviral action of the aqueous extract of propolis was tested against Chikungunya. The experimental procedures were the same as mentioned above, only with a greater amount of virus (1000 TCID/50%) and 5% propolis. As can be seen in Figure 7, a 256-fold reduction in viral replication was observed when crude propolis was added or a 512-fold reduction when the purified fraction was used.

**Figure 7:**
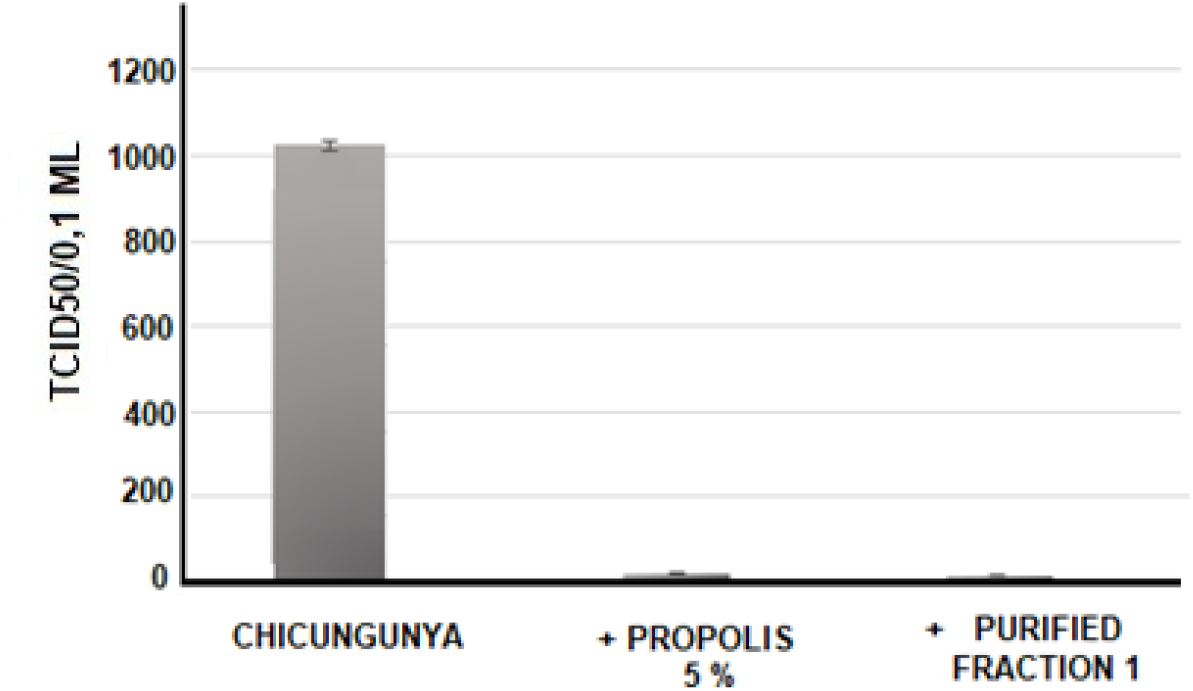
Antiviral activity of the aqueous extract of propolis from *Scaptotrigona* aff*. postica* (5% v/v) against the Chikungunya virus. The propolis extract or the purified fractions were added to the VERO cell cultures. 1 hour later, serial dilutions (r:2) of the Chikungunya virus (initial titer of 1000 TCID 50/0.1ml) were added to the culture. The culture was maintained at 37 °C for 72 hours and the culture was observed daily to determine the cytopathic effect. The final titer was determined by the highest dilution of the virus to cause a cytopathic effect in the culture. The data in the figure is representative of 4 experiments (performed in duplicate each)

In the photos in Figure 8, it can be seen that there was a clear reduction in the cytopathic effect caused by 5% (v/v) of propolis or its purified fraction against the Chikungunya virus (72 hours after infection).

**Figure 08.**
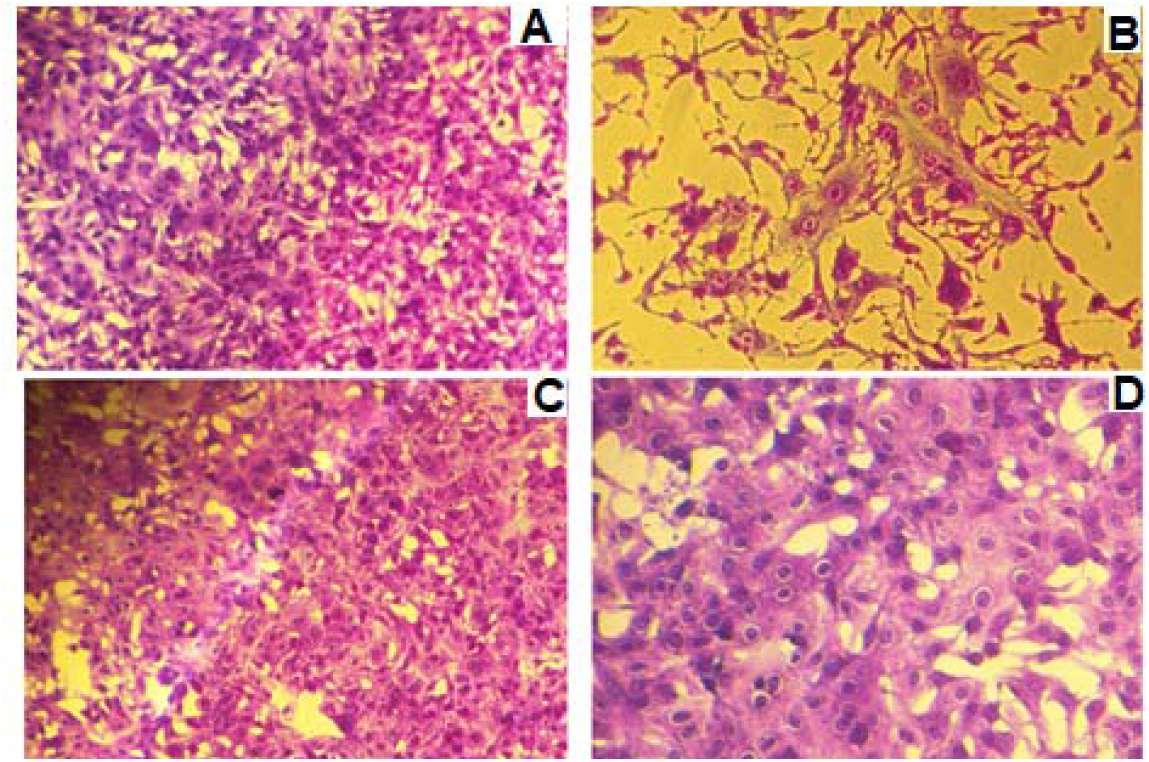
Photomicrography of normal VERO cell cultures (A), infected with Chikungunya virus (B), and infected cells treated with 5% v/v propolis from *Scaptotrigona* aff. *postica* (C) or with infected and treated with 5% of purified fraction of propolis (D). The cultures were kept for 3 days at 37°C. Cells were observed daily under an inverted microscope at 10x magnification (A, B, and C) and 20x in photo D. After this period, the culture supernatant was removed and the cells stained with crystal violet 025%.

#### 3.2.2. Effect of the concentration of propolis aqueous extract on viral replication

The action of concentration of propolis aqueous extract on antiviral activity was tested with Zicavirus or with Mayaro. For this, the VERO cell cultures were treated with 2, 5, or 10% propolis and after 1 hour the cells were infected with 128 or 512 TCID/50% of Zica virus or Mayaro virus. The cultures were observed for 72 hours and the viral titer was defined as the highest dilution of the virus capable of inducing a cytopathic effect in the cells. As can be seen in Figure 9, the antiviral action of propolis is dose-dependent, and the viral reduction with the concentration of 10% of propolis extract was 64-fold to Zicavirus and 128-fold to the Mayaro virus (99,6% of reduction).

**Figure 9:**
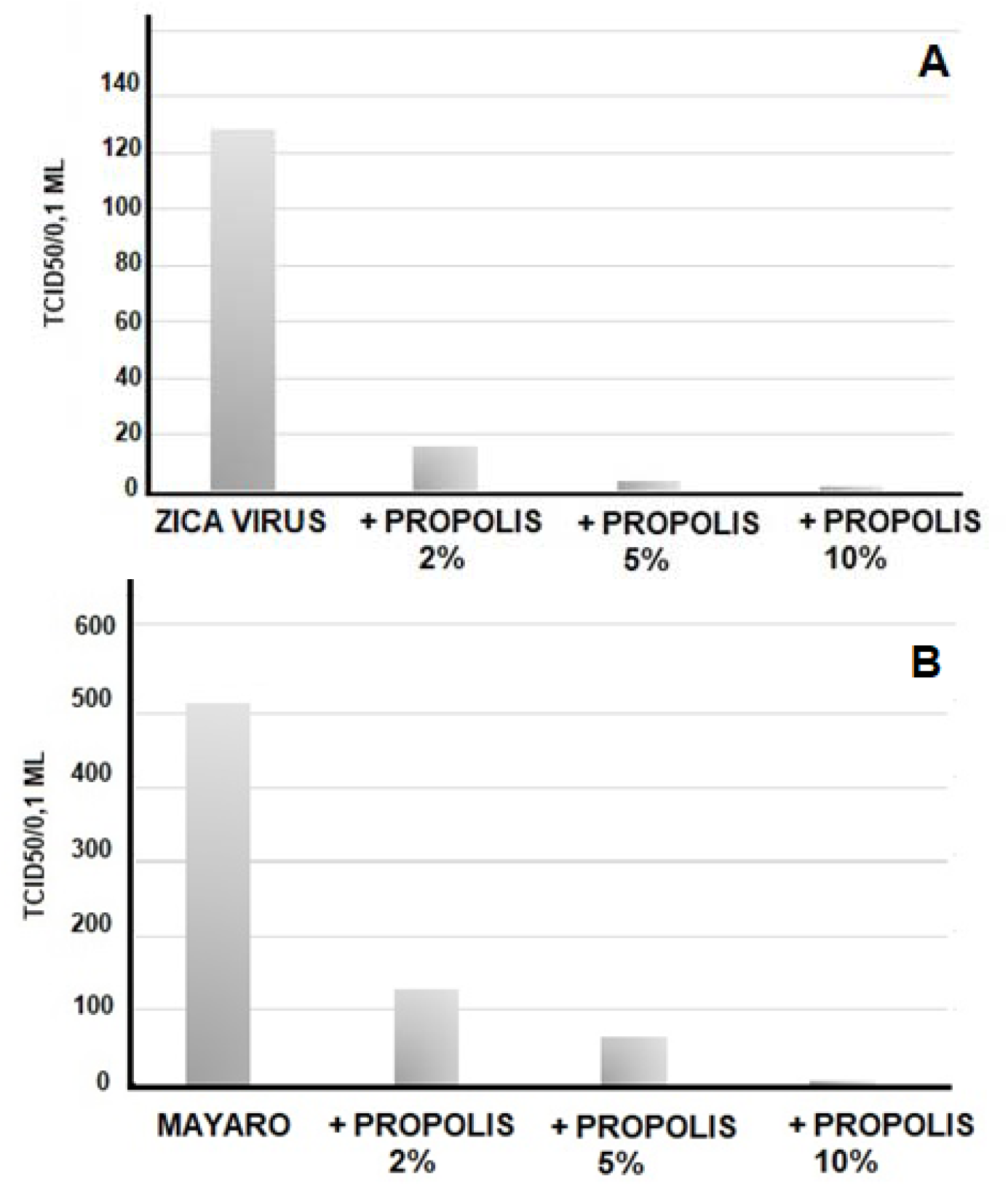
Antiviral action of the concentration of the aqueous propolis extract of *Scaptotrigona* aff. *postica* against Zicavirus (A) or Mayaro virus (B). Aqueous propolis extract (2, 5, or 10% v/v) was added to the VERO cell culture. 1 hour after this procedure, serial dilutions (r:2) of Zicavirus or Mayaro virus (initial titer of 128-512 TCID 50/0.1ml) were added to the culture. The culture was maintained at 37 °C for 72 hours and the culture was observed daily to determine the cytopathic effect. The final titer was determined by the highest dilution of the virus to cause a cytopathic effect in the culture

#### 3.2.3. Effect of time of addition of aqueous propolis extract on viral replication

The effect of the time of addition of the aqueous extract of propolis on the antiviral activity was tested against different dilutions of Zicavirus (from 0 to 2048 TCID/50/0.1 ml). For this, VERO cell cultures were treated with 5% propolis 2 hours before infection, 2 hours after infection, or treated with a mixture of virus + propolis kept together for 2 hours. The culture was observed for 6 days and the viral titer was defined as the highest dilution of the virus capable of inducing a cytopathic effect in the cells. As can be seen in Figure 10, the antiviral action of propolis was more effective when propolis was added 2 hours after infection. The experiments were performed in triplicate.

**Figure 10:**
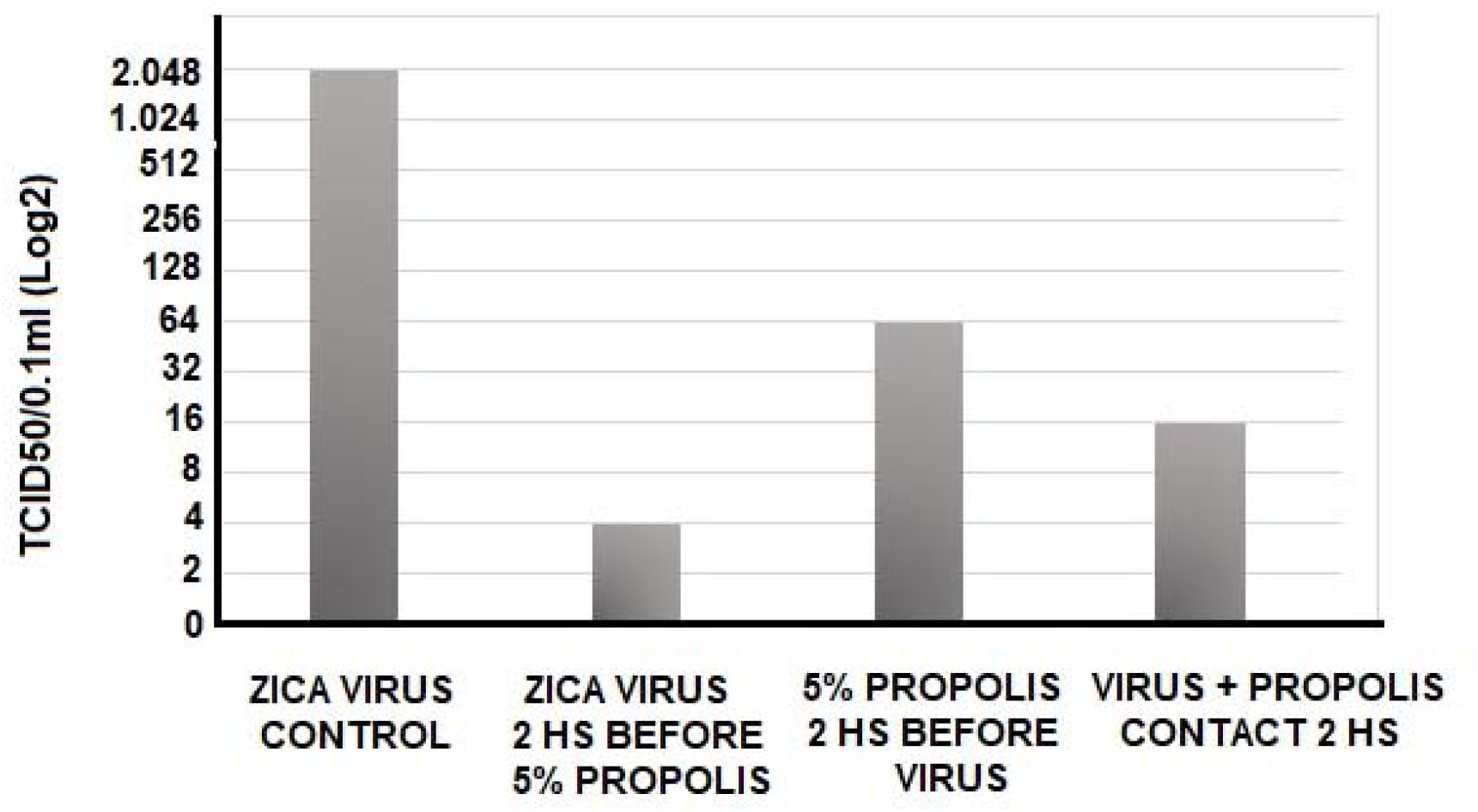
Action of propolis aqueous extract (TOI) addition time on antiviral activity against different Zicavirus dilutions (from 0 to 2048 TCID/50/0.1 ml). For this, VERO cell cultures were treated with 5% propolis 2 hours before infection, 2 hours after infection, or treated with a mixture of virus + propolis kept together for 2 hours. The culture was observed for 6 days and the viral titer was defined as the highest dilution of the virus capable of inducing a cytopathic effect in the cells.

## DISCUSSION

Arboviruses are viruses transmitted by arthropods (Arthropod-borne virus) and are so named because part of their replicative cycle occurs in insects. They are transmitted to humans and other animals by the bite of blood-sucking arthropods (mosquitoes). Of the more than 545 known arbovirus species, about 150 cause disease in humans. (3). Arboviruses have represented a major challenge to public health, due to climate and environmental changes and deforestation that favor amplification, and viral transmission, in addition to the transposition of the barrier between species. Among the arboviruses that cause diseases in humans and other warm-blooded animals are Dengue, Zica, Chikungunya, and Mayaro, among many others. These viruses are a major problem for which there is no specific treatment Treatment is usually focused only on relieving symptoms because there are no drugs that specifically attack the virus (53). In general, medications are indicated to control fever, headache, skin spots, and body pain, the same used in patients with dengue, zikavirus, or other arboviruses. That is why it is extremely important to develop a drug that acts directly on the virus and not just on the symptoms. Various studies have reported the antiviral activity in products obtained from invertebrates and their products. Among the various animal sources of antiviral studies, propolis has gained great attention for obtaining new drugs (54, 15), mainly because of its extensive and varied chemical composition (15, 29, 55) that generate huge amounts of molecules of high pharmacological value. Peter et al (56), describe the antiviral effect of three hydroalcoholic extracts of propolis (brown, green, and jataí bees (Tetragonisca angustula), against bovine herpesvirus type-1 (BoHV-1) and bovine viral diarrhea Virus (BVDV). This propolis has shown high efficacy in both cellular treatments (post and pre-exposure) against BoHV1. Extracts from Jataí showed activity agains46, t both viruses (BoHV-1 and BVDV) infection in the pre-test, whereas brown propolis demonstrated action only against the BoHV-1 in the pre-infection method., being that, the green propolis, led to a reduction in viral titer of 4.33log, while was observed a reduction of 3.5log to brown propolis and of the 3.24log to jataí propolis. Like this one, to date, most of the scientific work on the therapeutic activities of propolis has been carried out with the bee species Apis mellifera. However, some studies using propolis from stingless bees have been described, demonstrating its therapeutic potential against viruses of importance in human and veterinary health such as human herpes virus type 1 (46, 57, 58), Herpes Vírus Bovino (BoHV-5) (59), Parvovírus Suíno (PPV) (60), vírus da imunodeficiência human (HIV) (61) and influenza virus (62).

One of the advantages of propolis as a therapeutic agent is, in general, the absence of toxicity to humans and animals (14). In the literature, studies have been observed in which the concentrations of propolis used in antiviral assays vary from 0.0003% to 5% (57), with no morphological alterations being observed in the cells (14). However, Peter et al.(56), observed cytotoxicity of 0.097µg/mL for brown and green propolis and 0.39µg/mL for jataí propolis. In a similar study, Bankova et al.(15) evaluated the antiviral activity of a hydroalcoholic extract of Canadian propolis, where it was observed that the maximum tolerated non-cytotoxic concentration of propolis in MDBK cells was 0.032mg/mL. This relative toxicity may be related to the extraction method used, which was a hydroalcoholic extraction, which removes a good amount of bioactive substances present in propolis. Contrary to what is normally used, our experiments were carried out with aqueous extraction. With this procedure, substances that are not isolated in alcoholic extracts are obtained, which leads to the discovery of new molecules with therapeutic potential, in addition to avoiding the presence of other substances that may be toxic in addition to those present in the material under study. In our study, with the aqueous extraction, we did not observe any toxic effect on VERO cells, when we used final concentrations of up to 10% of propolis, which allows for obtaining a higher concentration of bioactive products in the formulation.

There are several reports of the antiviral activity of propolis. However, the mechanisms of action of propolis on viruses are not yet fully elucidated, but some studies seek to identify where this activity occurs. Bankova et al. (15) evaluated an alcoholic extract of propolis against Herpes simplex type 2 (HSV2) before contact with the cell, and suggest that the action of propolis occurs in the structure of the viral envelope or by modifying structural components necessary for adsorption or entry of the virus into the cell. The same study was carried out against Herpes simplex type 1 (HSV1), and the data suggest that when propolis is added before or at the time of infection, a greater virus-inhibiting effect is observed. In the study by Peter et al (56) with extracts of brown propolis and jataí, a greater effect of propolis was observed when previously inoculated the cells, before viral infection, suggesting the destruction of the virus inside the cell. However, the same result was not observed when this propolis sample was inoculated previously to BVDV, possibly showing different mechanisms of viral neutralization. There appears to be a wide variation in the activity of different types of propolis in antiviral activity, as when Peter et al (56) used green propolis extract, contrary to what was observed with brown and Jatai propolis, they did not observe significant antiviral activity in any of the experiments used. These data with green propolis differ from those obtained by Fischer et al. (59), who evaluated the effect of the antiviral activity of a Brazilian green propolis sample against bovine herpesvirus type 1 and BVDV. These differences may be related to the chemical composition of propolis from different crops. Recently, in a study carried out by us with 12-month samples of propolis from *Scaptotrigona* aff. *postica*, we have observed a great variation in the chemical composition of propolis according to the harvest month. This variation in chemical composition is related to the variation of flowering plants in each month, which bees use for the production of honey and propolis. Unlike other studies carried out with propolis from other sources, which derives exclusively or mostly from a single botanical source, the propolis from Barra do Corda used in our studies is composed of several sources, as it was obtained in a region of native forest, with a very large variety of plants (29).

One of the positive aspects of this propolis is that its origin is in a native region, without the use of pesticides, which prevents its contamination by other unnatural products. Cueto et al. (63) also reported the antiviral action of two ethanolic extracts of propolis from Rio Grande do Sul (Brazil), against some animal viruses, such as feline Calicivirus, Canine Adenovirus type 2, and Bovine Viral Diarrhea Virus. In this study, the presence of flavonoids such as rutin, quercetin, and gallic acid was identified. In this work, antiviral activity against BVDV and CAV-2 was observed when propolis was added before viral infection. In a study carried out by us, (46) we determined the antiviral action of a hydroethanolic extract of *Scaptotrigona* aff. *postica* from the region of Barra de Corda, state of Maranhão, Brazil against Herpes Simplex Virus (HSV-1). The chemical analysis of propolis showed higher amounts of pyrrolizidine alkaloids and C-glycosyl flavones, which is a compound detected for the first time in a sample of origin of meliponian bees. Apparently, the viral type also influences the results of antiviral activity. Peter et al (56), observed that jataí propolis - T. angustula) obtained better antiviral results against RNA virus (BVDV) than when using DNA virus (BoHV-1). The BVDV genome consists of a single strand of RNA, surrounded by a lipid envelope, which constitutes a structural difference concerning BoHV-1, which is a DNA virus (63). Eventually, this may explain the greater action of the propolis obtained from the jataí bee. Yildirim et al. (58) observed a significant reduction in HSV-1 and HSV-2 replication with Jatay propolis. However, toxic action was observed at concentrations of 200 and 400 μg/mL. The time of action of propolis also varied according to the virus used. While the replication of HSV-1 started 24 h after infection, for HSV-2 the start of replication was after 48 h of incubation. In our study, propolis of *Scaptotrigona* aff *postica* was tested, for potential antiviral activity against arboviruses. Experiments with total propolis led to an average 256-fold reduction in replication of the three tested viruses, while the semi-purified protein led to a 512-fold reduction in Chikungunya virus production, a 64-fold reduction in Mayaro replication, and a 16-fold reduction in Zica virus production. The action of propolis was higher when the cell culture was infected 2 hours before the addition of propolis, which suggests an intracellular mechanism of action and that the protein may act as a constitutive agent that affects the innate antiviral immune response. IFNs, cytokines and chemokines are seen as essential for efficient host defense against infection. Melchjorsen et al (64) showed that the initial response of HSV-infected cells is the secretion of antiviral substances such as defensins and nitric oxide and cytokine production, including IFN and chemokines. Earlier we also reported that type I IFNs stimulate Nitric Oxide Production and Resistance to *Trypanosoma cruzi* Infection (65). As occur with trypanosomes, we believe that the mechanism of action of some antivirals is to stimulate cells to produce interferon, nitric oxide, or other cytokines. Nevertheless, its mechanism of action needs further investigation. Concluding, this study showed antiviral activity against various arbovirus. It is can be an important target for the development of new compounds as an alternative to commercial antivirals. However, in addition to the antiviral activity, it is expected to obtain many other pharmacological activities in the propolis of *Scaptotrigona* aff. *postica*, given the great complexity in the composition of this product and the great variability in the seasonal chemical composition that we observed in this propolis (29)

## Supporting information

Propolis

## Acknowledgment

Laboratory of Parasitology and Protein Chemistry Laboratory Team (Butantan Institute).

